# Molecular mechanism of substrate transport by human peroxisomal ABCD3

**DOI:** 10.1101/2025.05.21.655323

**Authors:** Meghna Gupta, Nitesh Kumar Khandelwal, Devin J. Seka, Sree Ganesh Balasubramani, Miles Sasha Dickinson, Alexander Myasnikov, Ignacia Echeverria, Robert M. Stroud

## Abstract

Peroxisomes are eukaryotic oxidative organelles involved in numerous metabolic functions that include fatty acid oxidation, bile acid synthesis, and detoxification of reactive oxygen species. ATP-binding cassette transporters of the D subfamily (ABCD1-3) mediate the import of CoA thioesters of fatty acids into the peroxisome. ABCD3, the most abundant of these transporters in the peroxisomal membrane, facilitates the transport of a broad spectrum of substrates including branched-chain fatty acids, very long-chain fatty acids, bile salt intermediates, and dicarboxylic acids. Mutations in ABCD3 are associated with defects in congenital bile acid synthesis and variants of Zellweger syndrome. The structural and functional details of the human ABCD3 transporter remain unclear, despite its significance. In this study, we report the cryogenic sample electron microscopy (cryo-EM) structures of full-length human ABCD3 in its apo state and bound to one of the physiological substrates (phytanoyl-CoA) at resolutions of 3.33 Å and 3.13 Å, respectively. Our biochemical assays reveal that substrate binding induces ATPase activity in ABCD3, suggesting a substrate-dependent conformational change. Structural comparison of the apo and substrate bound states demonstrate that the substrate interaction brings nucleotide-binding domains closer, providing a mechanistic basis of substrate induced ATPase activity. These findings offer critical insights into the transport mechanism of ABCD3 and lay a structural foundation for understanding its role in peroxisomal metabolite import and related diseases.

**Significance Statement:** Peroxisomes are involved in essential cellular metabolic processes that include fatty acid oxidation, bile acid synthesis, and detoxification. The ABCD subfamily of membrane transporter proteins transport fatty acyl-CoA molecules from the cytosol into peroxisomes. Among the ABCD subfamily members, ABCD3 transports branched chain fatty acids and a range of other important metabolites. Dysfunction in ABCD3 has been associated with several inherited and acquired diseases. Using cryo-EM, we determined high-resolution structures of full-length human ABCD3 in both unbound and phytanoyl-CoA-bound forms. We show that substrate binding stimulates ATP hydrolysis activity by apposition of the two nucleotide binding domains of ABCD3, adding structural insight into the mechanism of action of ABCD3 and its role in peroxisomal function and disease.

## Introduction

Peroxisomes are organelles in all eukaryotic cells that harbor more than fifty enzymes, and membrane proteins that balance oxidative stress, mediate α- and β-oxidation of fatty acids, synthesize bile acids, catabolize D-amino acids and polyamines, and biosynthesize cholesterol, docosahexaenoic acid, and plasmalogens (1-4). Coenzyme A (CoA) thioesters of fatty acids are imported into peroxisomes by members of the D family of ATP-binding cassette (ABC) transporters for fatty acid oxidation (5). ABC transporters use the energy of ATP hydrolysis to transport substrates across the membrane (6, 7). Of the four human ABC subfamily D transporters, ABCD1-3 localize to the peroxisomal membrane and are involved in transport of acyl-CoA substrates (8-10). Human ABCD transporters function as homodimeric complexes, with each protomer consisting of a transmembrane (TMD) and a nucleotide-binding domain (NBD). These half-transporters form obligate dimers to make a fully functional transporter (9). ABCD3 protein has 37.65% and 39.15% sequence identity with ABCD1 and ABCD2, respectively. ABCD1 and ABCD2 share 64.65% identity with each other and have an overlap in substrate preference, constituting C18-C24 fatty-acyl CoA (4, 10). ABCD3 is one of the most abundant proteins on the peroxisomal membrane and transports diverse substrates-branched-chain fatty acids, long chain fatty acids, bile salt intermediates-dihydroxycholestanoyl-CoA and trihydroxycholestanoyl-CoA, dicarboxylic acids (11, 12). Loss or dysfunction of ABCD3 disrupts metabolic pathways, causing congenital bile acid synthesis defect 5, and certain types of Zellweger syndrome (13, 14). Alteration in expression levels of ABCD3 is associated with intrahepatic cholestasis, oculopharyngodistal myopathy, and its higher expression is correlated with better prognosis in various cancers (15-19).

Recently, Li et.al. reported that they attempted to overexpress and purify full-length human ABCD3 from HEK293F cells but were unable to obtain enough for structural characterization (20). Due to this limitation, they employed a chimeric version of the ABCD3 transporter to successfully determine its structures in both substrate-bound and ATP-bound conformations (20). However, the structure of the full-length human ABCD3, particularly in the apo state, remains undetermined, reflecting a significant gap in our understanding. Structural insights into ABCD3 in its various conformational states are essential for elucidating the complete mechanism of its transport cycle and the conformational transitions it undergoes during substrate translocation.

In the present study, we determined structures of detergent extracted human ABCD3 transporter using single particle cryo-EM for apo, and substrate-bound states at 3.33 Å and 3.13 Å resolution, respectively. Our ATPase activity experiments showed that purified ABCD3 is active at a basal level and exhibits an increase in activity with substrates phytanoyl-CoA, and palmitoyl-CoA. Structural comparison between the apo and phytanoyl-CoA-bound states of ABCD3 shows that substrate binding promotes the dimerization of the nucleotide-binding domains (NBDs), providing a mechanistic explanation for the observed substrate-induced ATPase activity. In summary, our integrated approach using cryo-EM, biochemical assays, and atomistic molecular dynamics (MD) simulation provides a comprehensive view of substrate recognition and conformational transitions, delineating key steps in the transport cycle of the peroxisomal fatty acyl CoA transporter-human ABCD3.

## Results

### Biochemical characterization and structure determination of human ABCD3

Human ABCD3 is a homodimer of ∼75 kDa. subunits, each half transporter having six transmembrane helices (TMHs) and one nucleotide binding domain (NBD). It is a half transporter like other ABCD subfamily ABC transporters and must form a dimer to function. Purification of full-length human peroxisomal ABC transporters has been challenging, and other approaches resorted to using a chimeric version of the protein (20, 21). We cloned the full-length human ABCD3 (hABCD3) and expressed it in *Sf9* cells. We successfully purified hABCD3 with high purity and obtained an amount adequate for structural and biochemical characterization (Fig.1A). We tested the functionality of detergent extracted and purified hABCD3 protein using ATPase assay and found it is active. The ATPase activity of hABCD3 is induced ∼5-fold in the presence of its substrates phytanoyl-CoA, or palmitoyl-CoA (Fig.1B).

**Fig. 1.**
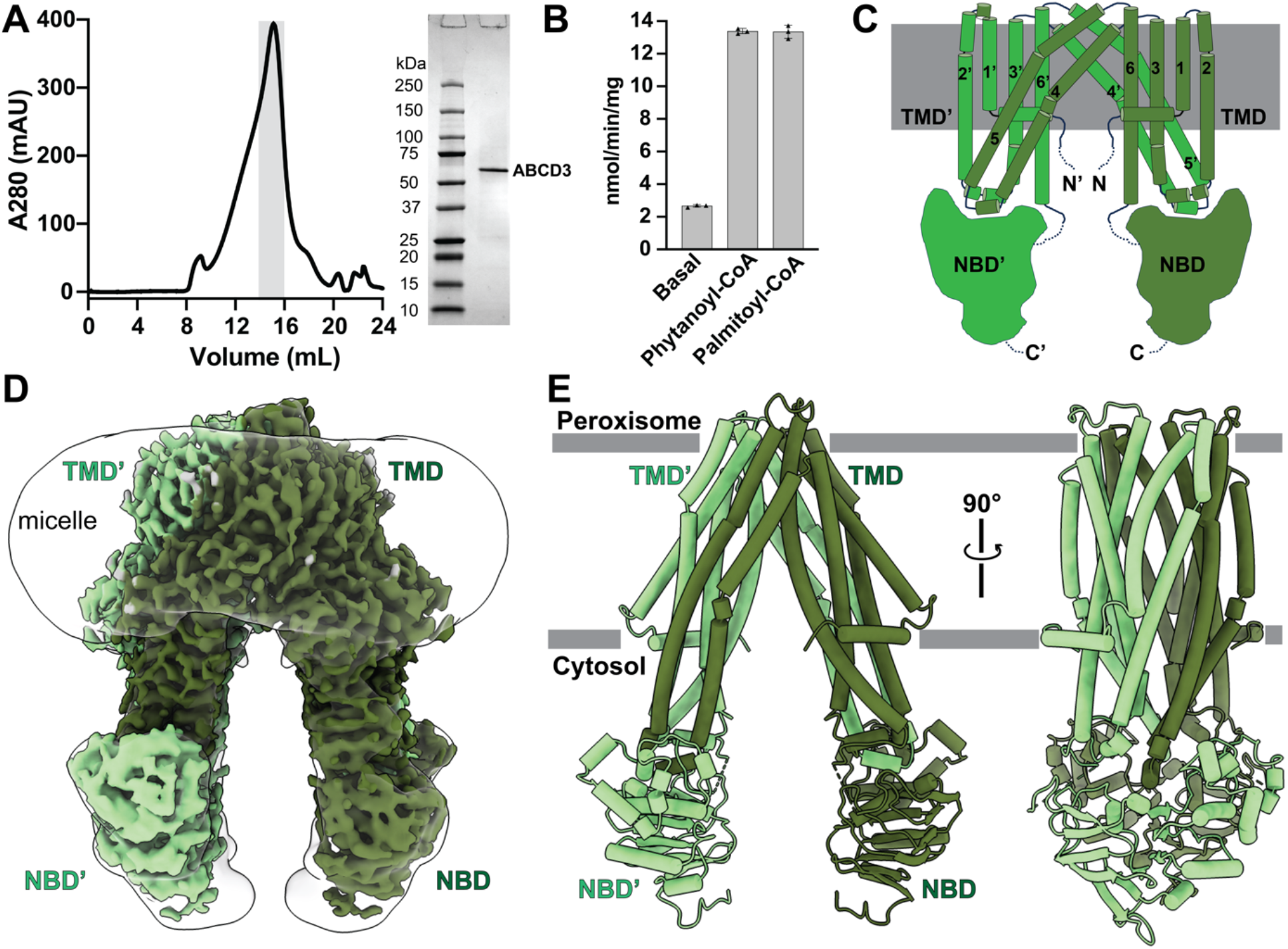
Human ABCD3 overall structure. **(A)** Size-exclusion chromatography profile of purified hABCD3 and SDS gel of peak fraction. **(B)** ATPase activity of ABCD3 at basal level and in presence of its substrates (phytanoyl-CoA and palmitoyl-CoA). **(C)** Overall topology of ABCD3. **(D)** Cryo-EM map of ABCD3 as homodimer in GDN detergent (dark green representing protomer A and light green representing protomer B). **(E)** Model of ABCD3 with color theme as shown in (D).

For structural studies, we prepared cryo-EM grids with hABCD3 samples and collected cryo-EM data as described in the Materials and Methods section. We processed the cryo-EM data and determined the hABCD3 apo structure at 3.33 Å resolution (Fig.1C-E and Appendix, Fig. S1). The hABCD3 apo structure was obtained in an inward facing conformation similar to the human ABCD1 transporter structures (21-24). The overall map density was sufficient to confidently model all six TMHs and NBD in both protomers (Appendix, Fig. S2), indicating that hABCD3 assembles as a two-fold symmetric homodimer. The hABCD3 TMHs are arranged in type IV ABC transporter topology where two TMDs each are swapped across the interface of the two halves (Fig. 1C). Here, TMH 4 and 5 of each protomer make transmembrane bundle with TMH 1, 2, 3, and 6 of the second protomer and *vice versa* (Fig. 1C and Fig. 1E). In this arrangement the intracellular loop (ICL) between TMH4 and 5 of one protomer is sandwiched between TMHs and NBD of the second protomer (Fig. 1C and Fig. 1E).

### Phytanoyl-CoA binding analysis with human ABCD3

Phytanoyl-CoA is a known substrate of hABCD3, and its transport from the cytosol into peroxisomes is essential for the α-oxidation of phytanic acid (25). We investigated the interaction between purified hABCD3 and phytanoyl-CoA using microscale thermophoresis (MST), which revealed a binding affinity of 54.9 ± 9.03 µM (Fig.2A). To explore the structural basis of this interaction, we prepared a complex of hABCD3 with phytanoyl-CoA for cryo-EM structure determination and resolved it to a resolution of 3.13 Å. Our structure reveals two phytanoyl-CoA molecules bound to the ABCD3 homodimer (Fig. 2B-C). We propose that the binding of phytanoyl-CoA induces a more compact conformation of the protein, contributing to the improved resolution (3.13 Å) of the ligand-bound state (Appendix Fig. S3) compared to the apo state (3.33 Å) (Appendix Fig. S1).

**Fig. 2.**
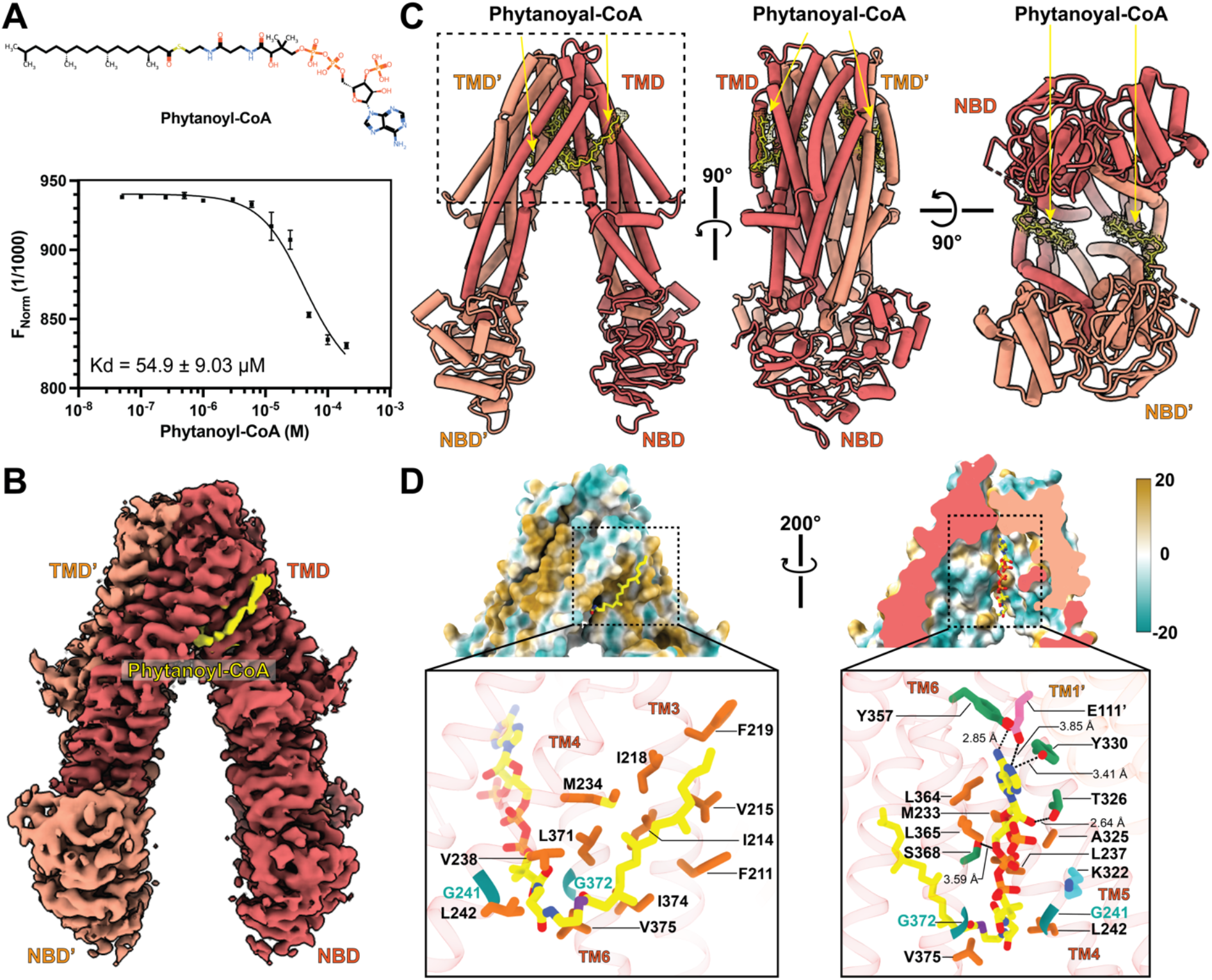
Phytanoyl-CoA bound Human ABCD3. **(A)** The binding affinity of phytanoyl-CoA for human ABCD3 determined using MST to be 54.9 ± 9.03 µM (n = 3) with error bar representing S.D. **(B)** Cryo-EM map of phytanoyl-CoA bound ABCD3. Density correspond to phytanoyl-CoA is yellow in color, dark orange color representing protomer A and light orange color representing protomer B. **(C)** Model of phytanoyl-CoA bound ABCD3 with color code as in (B). The density corresponding to the phytanoyl-CoA is shown in mesh. Left panel showing the view as B, middle panel rotation of left panel at 90° on y-axis. Right panel rotation of middle panel at 90° on x-axis. **(D)** Hydrophobicity surface view of ABCD3 around the box region shown in (C) showing the phytanoyl-CoA binding site. The lipid tail of phytanoyl-CoA outside the transmembrane helix cavity shown in upper left panel with insight of hydrophobic interaction network in lower left panel. The trimmed view of the ABCD3 substrate binding pocket with phytanoyl-CoA bound in it (upper right panel) with highlighting the interactions detail in lower right panel. Phytanoyl-CoA is colored in yellow, glycines flanking the cavity are in teal, hydrophobic residues are colored in orange, polar-uncharged residues colored in green, negatively charged residues are in pink, and positively charged amino residues are in cyan; colored by atom-oxygen (red), nitrogen (blue), and sulfur (violet).

In our phytanoyl-CoA bound ABCD3 map, we observed two extra densities between the TMDs, which we assigned to two phytanoyl-CoA molecules (Fig. 2B-C; Appendix, Fig. S4). Each molecule engages in a network of hydrogen bonds, hydrophobic contacts, and van der Waals interactions with TMHs 3-6 of one protomer and TMH1’ of the adjacent protomer (Fig. 2C-D). Phytanoyl-CoA is composed of hydrophilic coenzyme A and hydrophobic phytanic acid. Within the ABCD3 protein, this compound adopts a V-shaped conformation where its hydrophilic part (coenzyme A) is buried within the substrate binding pocket surrounded by transmembrane domains (Fig. 2D, right panel) and the hydrophobic phytanic acid tail comes out the pocket and runs upwards parallel to hydrophobic surface composed residues form TMH 3, 4 and 6 (Fig. 2D, left panel). The phytanoyl-CoA branched fatty acyl chain is tucked in the hydrophobic cleft formed from residues F219, I218, V215, I214, F211 of TMH3, M234, V238, L242 of TMH4 and L371, I374, V375 of TMH6 occupying 301.63 Å^2^ of surface area (Fig. 2D, left panel).

The CoA moiety makes multiple interactions within the hydrophilic pocket. The CoA adenine ring makes H-bonds with Y357 (TMH6) and Y330 (TMH5) and E111’ (TMH1’). The hydroxyl group present on the second carbon of the ribose ring makes hydrogen bond with T326 (TMH5) and the 3’-phosphate present on its ring is placed in adjacent proximity of K322 (TMH5) with the possibility of ionic interaction. The oxygen of first phosphate from the diphosphate interacts with S368 (TMH6) (Fig. 2D, lower right panel). At the bottom of the substrate binding cavity of the TMDs, two glycine residues G241 and G372 from TMH4 and TMH6, flank the pantothenic part of CoA allowing access of attached phytanoyl hydrophobic tail towards the bilayer (Fig. 2D).

### Molecular dynamics simulation reveals phytanoyl-CoA interacting residues

Phytanoyl-CoA is a large amphipathic substrate with distinct hydrophilic and hydrophobic moieties. We performed molecular dynamics (MD) simulations to track the interactions and conformational changes of phytanoyl-CoA within the ABCD3 substrate binding cavity, specifically investigating how its molecular size and amphipathicity influence dynamic behavior over time. An initial 50 ns equilibration was conducted with the ligand bound to the protein in a 1-palmitoyl-2-oleoyl-sn-glycero-3-phosphocholine (POPC) lipid bilayer membrane. Subsequently, three independent 500 ns production runs were generated to enhance conformational sampling, each initiated from a distinct starting configuration (Appendix, Fig. S5 and Fig. 3). During the simulation time across all three trajectories, phytanoyl-CoA stays within the cavity; however, it displayed significant conformational flexibility and variable interaction patters with different protein residues. We expect the long, hydrophobic lipid tail of phytanoyl-CoA, which is located outside the protein cavity (Fig. 2D), to be flexible. We focused on the CoA moiety of phytanoyl-CoA, which resides within the ABCD3 cavity. This moiety was divided into five parts (adenine ring, ribose-3-phosphate, diphosphate, pantothenate, and cysteine to examine the movement of the center of mass for each part individually (Fig. 3A-B). We observed that the free end of the substrate (adenine ring) and its pantothenate and cysteine near the exit point from the ABCD3 cavity, show greater movement compared to the central (ribose-3-phosphate, diphosphate). This is indicated by the standard deviations of the distances calculated based on the movement of the center of mass from its initial position (Fig. 3B). This flexibility leads to new interactions with nearby residues in the substrate binding cavity of ABCD3 (Fig. 3D). Specifically, phytanoyl-CoA in protomer A interacts with residues R373, R371, Q369, and K322 of protomer A, as well as R366’ and R373’ of protomer B within the cavity.

**Fig. 3.**
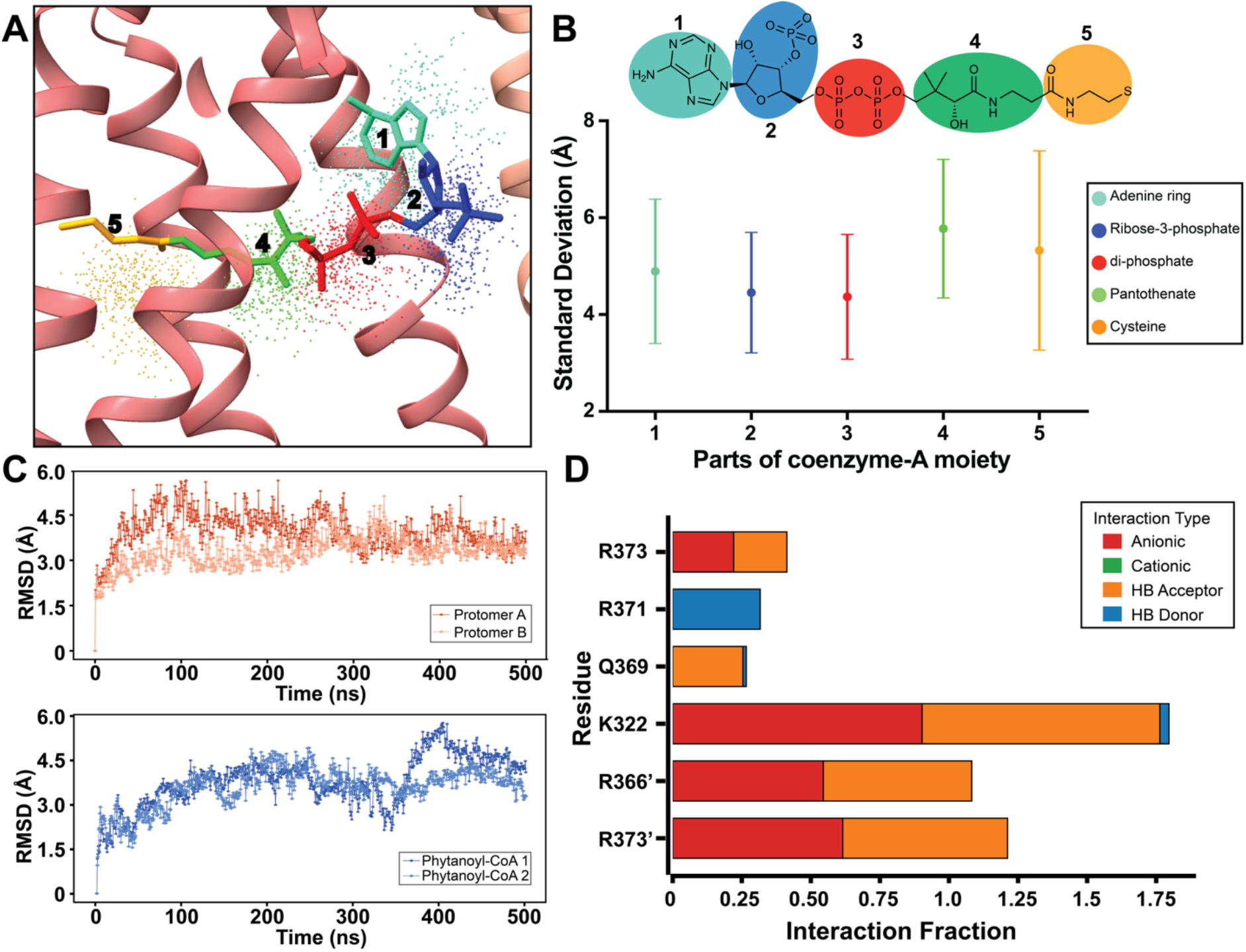
Molecular dynamics simulation of phytanoyl-CoA bound to hABCD3. **(A)** Structure and dynamics of Phytanoyl-CoA 1 within the dimer cavity. Its final conformation (sticks) is shown, segmented into adenine (Turquoise), ribose-3-P (Blue), diphosphate (Red), pantothenate (Green), and cysteine (Yellow). Correspondingly colored points illustrate the spatial volume explored by the center of mass (COM) for each segment during a representative trajectory. **(B)** Segmental flexibility of Phytanoyl-CoA 1. The standard deviation (SD, in Å) of COM displacement quantifies the positional fluctuation for each segment (colors as in A). Larger SD values for the terminal adenine and cysteine segments highlight their greater conformational flexibility. **(C)** System stability assessed by Root Mean Square Deviation (RMSD) relative to the initial structure, shown for a representative trajectory. Top: Protein backbone Cα RMSD for Protomer A (dark orange) and B (light orange). Bottom: Heavy atom RMSD for Phytanoyl-CoA 1 (dark blue) and 2 (light blue). **(D)** Persistent substrate-protein interactions identifying key binding residues. Interaction fractions (% time) depict significant hydrogen bonds and ionic contacts between Phytanoyl-CoA 1 and residues from Protomer A and B’ (prime denotes Protomer B). Displayed interactions represent consistent contacts, maintained for ≥10% of the time across all three replicate simulations.

### Substrate-induced changes in human ABCD3

Structural alignment indicates that both the apo and phytanoyl-CoA bound structures are in inward-facing open states with Root Mean Square Deviation (RMSD) of 1.362 Å between their respective Cα positions (Fig. 4). Phytanoyl-CoA interaction with ABCD3 brings the TMD of each half closer together such that the distance between E82s of TMH1 from both protomer reduced from 60.84 Å to 58.92 Å. This translates to larger movement between NBDs. The distance between V605 from NBD and R642’ from NBD’ reduced from 38.18 Å (Fig. 4A) to 34.28 Å (Fig. 4B).

**Fig. 4.**
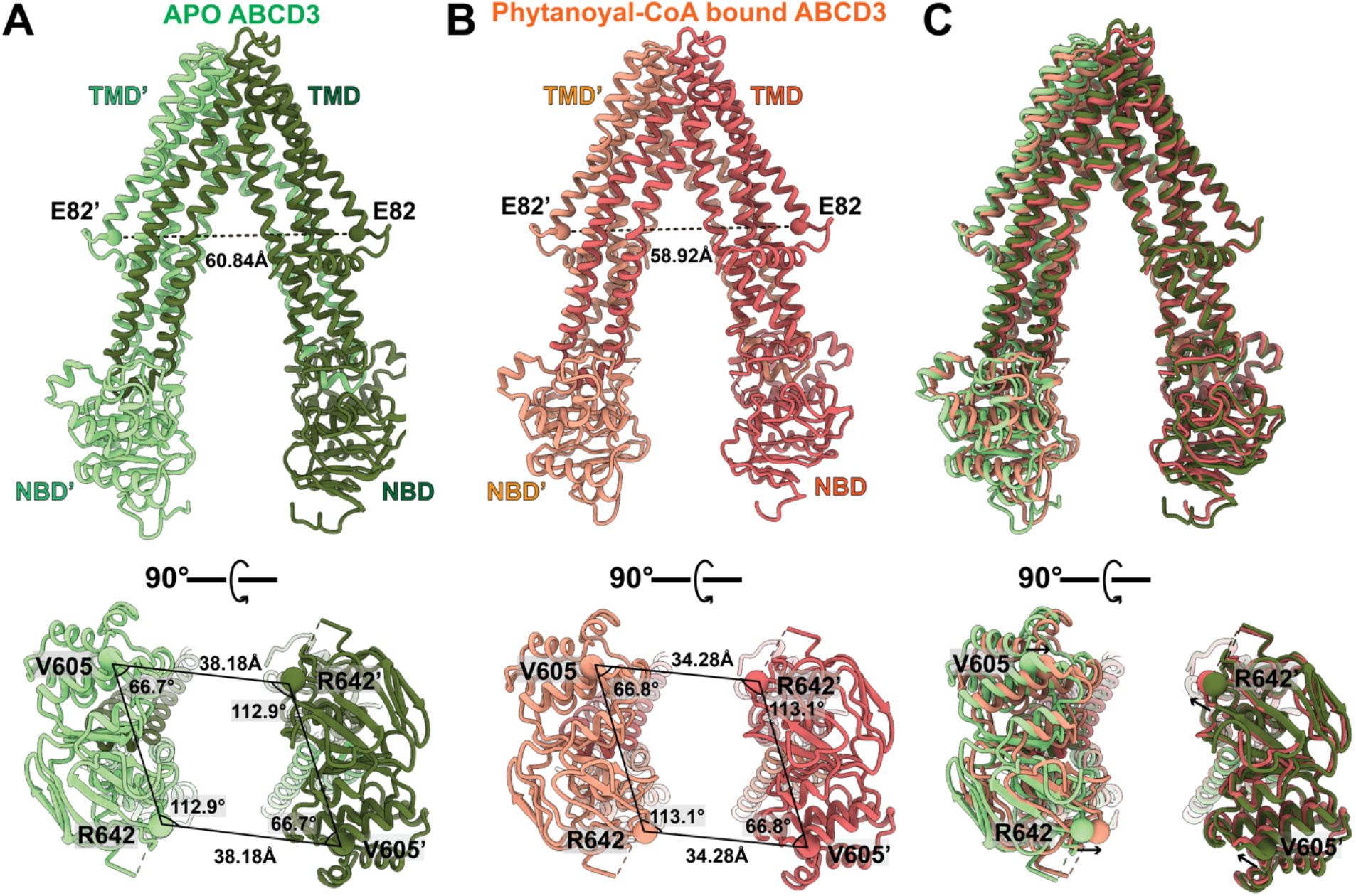
Phytanoyl-CoA binding induced changes in ABCD3. **(A)** Distance between TMDs and NBDs and angles between protomers in apo state **(B)** Phytanoyl-CoA bound state **(C)** overlay of apo and phytanoyl-CoA bound states. Change is measured as distance between E82 residue of TMH 1 of ABCD3 protomers **(A)** in apo state (top panel) to **(B)** in phytanoyl-CoA bound state (top panel). **(C)** overly of both structure (top panel). The change of NBDs position measured as angles and distance between the V605 and R642 residues of NBDs from each protomers of ABCD3. (A) in apo state (lower panel) to **(B)** in phytanoyl-CoA bound state (lower panel). **(C)** overlay of both structure (lower panel) highlighting the rearrangement in the relative positions of V605 and R642 residues of NBDs in apo (green sphere) and phytanoyl-CoA bound (orange sphere).

Phytanoyl-CoA interacts with TMH 3, 4, 5, 6 of one protomer and TMH1’ of second protomer. This interaction network pulls TMH4 and 5 closers to its cavity (Fig. 5A). This movement reduces the distance between phytanoyl-CoA and ABCD3 residues (Fig. 5B). We compared the distance between multiple ABCD3 residues in apo and substrate-bound states [Y330 (TMH5), T326 (TMH5), Y357 (TMH6), S368 (TMH6)] which make hydrogen bonds with phytanoyl-CoA. In the apo state, the distance between these residues is in the range of 4.1-

**Fig. 5.**
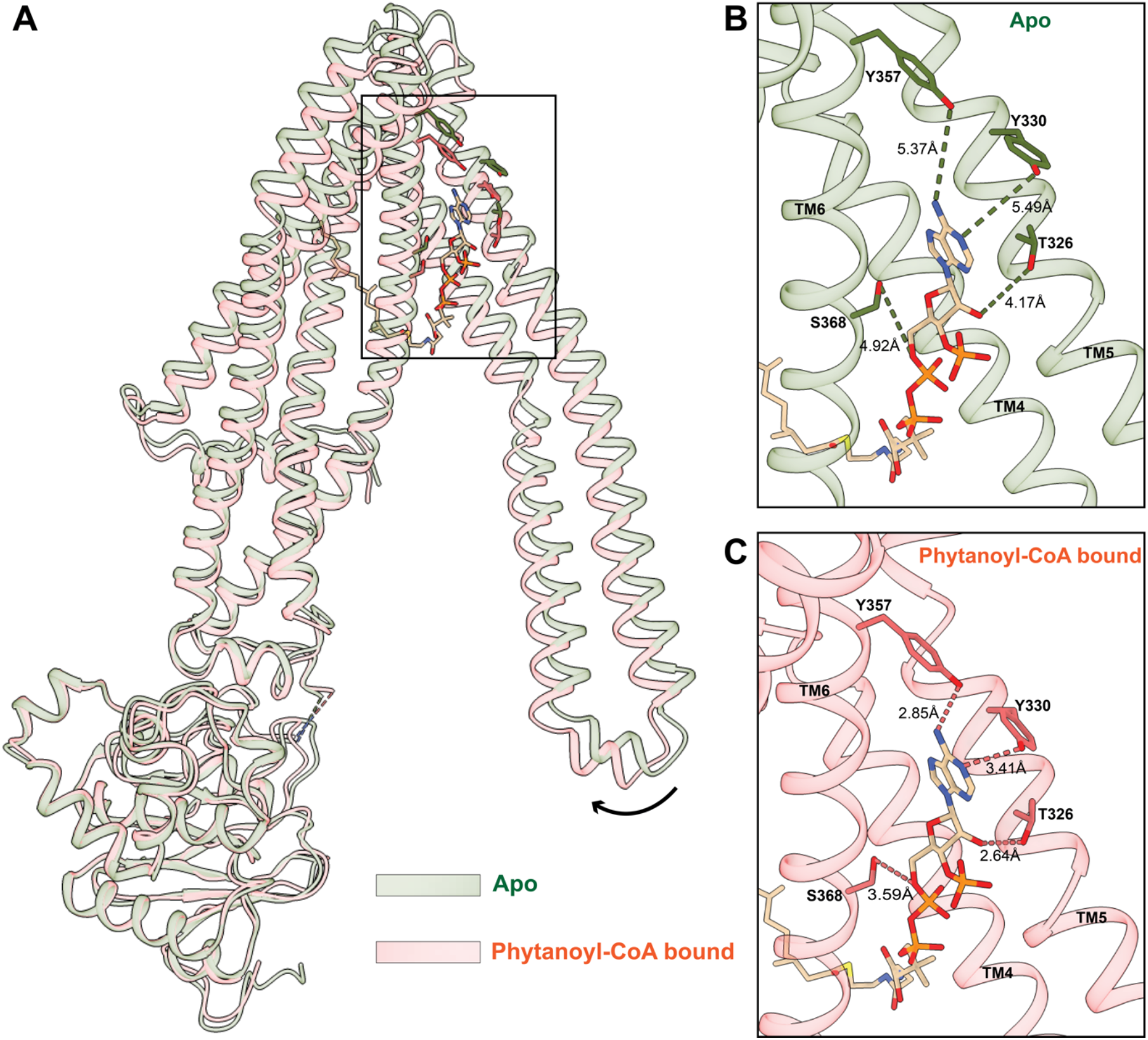
Comparison of apo and phytanoyl-CoA bound ABCD3 protomer. **(A)** Comparison of the transmembrane helices of apo and phytanoyl-CoA bound protomer A obtained from superimposition of the NBD region. In phytanoyl-CoA bound state, TMH 4 and 5 rotate and move inwards. **(B)** The rearrangement of TMH 4 and 5 in the apo state increases the distance between phytanoyl-CoA and residues (Y330, T326, of TM5 and Y357, S368 of TM6) (upper panel) from those in the phytanoyl-CoA-bound structure (lower panel). **(C)** Phytanoyl-CoA binding leads to a decrease in distance between phytanoyl-CoA and residues (Y330, T326, of TM5 and Y357, S368 of TM6) (lower panel) in comparison to the apo-state. 5.5 Å (Fig. 5B, upper panel), and this changes to a favorable range (2.6-3.6 Å) for interaction of the phytanoyl-CoA bound state (Fig. 5B, lower panel).

## Discussion

Overexpressing full-length human peroxisomal ABCD subfamily transporters (ABCD1– D3) has proven challenging, with several studies reporting insufficient yields of recombinant proteins for structural analysis. We successfully expressed and purified full-length human ABCD3 (hABCD3) from insect Sf9 cells. ATPase activity measurements confirmed that the purified protein was active and exhibited phytanoyl-CoA and palmitoyl-CoA induced ATPase activity (Fig. 1B). Our observed basal ATPase activity was low, which subsequently increased ∼5-fold upon introduction of its physiological substrates, demonstrating substrate dependent ATP hydrolysis. Similar observations were made by Chen et. al. in the case of another ABCD subfamily transporter (ABCD1) where basal ATPase activity was very low and increased in the presence of C22:0-CoA (21). Jia et. al., in another study on ABCD1 (26), report very high level of ATPase activity (∼180 fold higher than Chen et. al.) at the basal level. The major difference between these assays was that Chen et. al. used chimeric ABCD1 (first 63 amino acids at N-terminal of human ABCD1 were replaced by 65 amino acids of *Caenorhabditis elegans* PMP-4), and in contrast, Jia et. al. used N-terminal truncated ABCD1 (54 residues N-terminal deleted). In the same year, research from Le et. al. reported specific ATPase activity of full-length ABCD1 in the presence of different substrates in the range of ∼20 nmol/mg/min (22). This is comparable to the ATPase activity of full-length human ABCD3 in the presence of substrates we report here. However, the full-length ABCD1 ATPase activity reported by Le et. al. is very low in comparison to chimeric ABCD1 (21) as well as truncated ABCD1 (26).

In the case of ABCD3, higher ATPase activity was observed in chimeric ABCD3 (20). We reason that the N-terminal of human ABCD family members may play a regulatory role in ATPase activity, and hence replacing it with the *C. elegans* PMP4 sequence or deleting it results in elevated ATPase activity. This is supported by another recent publication where a comparative ATPase assay of full-length and N-terminal (54 residues) truncated ABCD1 showed higher activity in the case of the truncated version of ABCD1 (24).

In our phytanoyl-CoA-bound structure, we observed two additional densities compared to the apo structure corresponding to phytanoyl-CoA. MST-based interaction analysis between ABCD3 and phytanoyl-CoA revealed a Hill coefficient value of approximately 2.33, indicating strong cooperative binding. This interaction brings the two halves of ABCD3 closer together, leading to a reduced distance between the NBDs (Fig. 4). The substrate-dependent narrowing of the NBD positioning may facilitate faster dimerization upon ATP binding, potentially explaining the enhanced ATPase activity observed in the presence of the substrates (Fig. 1B).

Substrates for ABCD family transporters are mostly long carbon lipidic tail containing molecules. The flexible nature of these substrates facilitates interaction with different residues during the transport cycle. We assessed substrate binding to ABCD3, comparing our phytanoyl-CoA-bound structure with ABCD1 bound to oleoyl-CoA (23) and C22:0-CoA (21) (Appendix, Fig. S6). Superimposition of phytanoyl-CoA-bound ABCD3 with oleoyl Co-A bound or C22:0-CoA ABCD1 shows different substrate binding positions (Appendix, Fig. S6B). Although the CoA moiety is common in all these substrates the C22:0-CoA CoA interaction spans from one TMD across to the other TMD′ of the dimer (Appendix, Fig. S6B). CoA of oleoyl-CoA is placed in the ABCD1 transporter pocket comparable to the phytanoyl-CoA-bound ABCD3 (Appendix, Fig. S6A). In ABCD1, for both oleoyl-CoA and C22:0-CoA, the fatty-acyl chain is buried between the TMDs while the fatty-acyl chain in phytanoyl-CoA-bound ABCD3 is present outside of the TMDs. In the ABCD1 structure with C26:0-CoA, fatty-acyl chain is present outside of TMDs like our substrate-bound ABCD3 (24). One possibility justifying the differences in substrate-binding is that these structures represent distinct stages of the transport cycle. A chimeric ABCD3 protein structure with phytanoyl-CoA state was published in an inward-facing state during our manuscript preparation. The overall structure is very similar with an RMSD of 0.973 Å across 567 Cα atoms, with a slight difference in the binding poses of phytanoyl-CoA (Appendix, Fig. S7).

Structural analysis of ABCD3 in two distinct states (apo and substrate-bound), together with previously published ATP-bound states, allowed us to propose a model for ABCD3-mediated phytanoyl-CoA transport (Fig. 6). In its resting state, ABCD3 exhibits an inward-facing conformation where the two nucleotide-binding domains (NBDs) remain apart. Upon cooperative substrate binding, an induced fit causes the transmembrane domains (TMDs) to shift closer together, forming a narrower inward-facing transport cavity. This conformational change may facilitate the dimerization of the NBDs upon ATP binding, transitioning ABCD3 to an outward-facing conformation. This shift from inward-facing to outward-facing conformation enables the release of phytanoyl-CoA across the membrane completing the transport process. The substrate may be released in its intact form or hydrolyzed into phytanic acid and CoA via potential ABCD3 thioesterase activity. Finally, ATP hydrolysis and subsequent phosphate release trigger conformational changes that reset ABCD3 to its resting state, ready for another transport cycle.

**Fig. 6.**
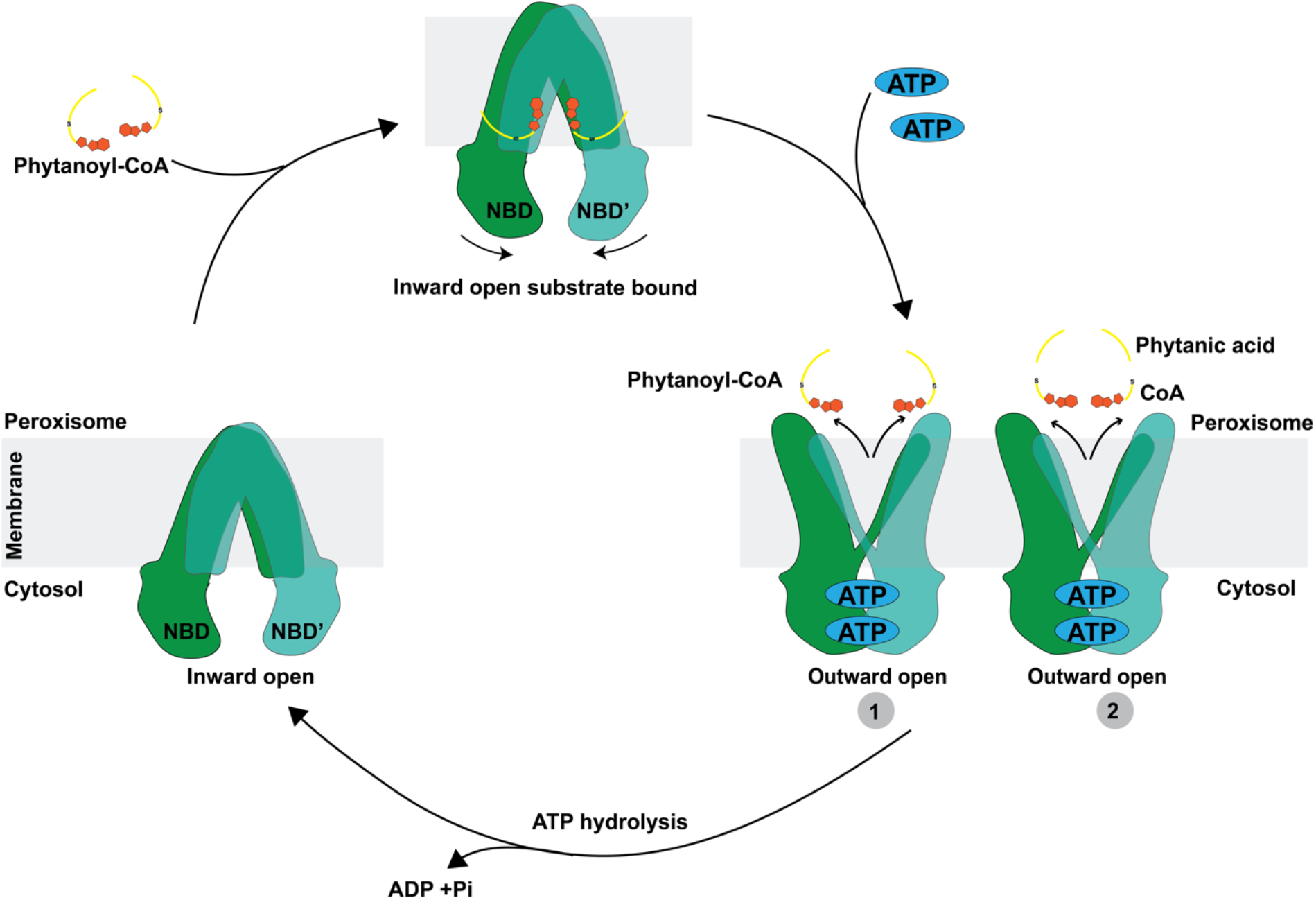
Proposed model of the transport cycle of ABCD3. Substrate (phytanoyl-CoA) binds to inward-open state of ABCD3 and brings NBDs closer together. ATP binding brings NBDs further together resulting in an outward-open state (PDBID:8Z0F) (20). In this state, substrate-release could happen in two possible ways. Phytanoyl-CoA can either be released to the peroxisome as an intact molecule or ABCD3 can act as a thioesterase resulting in release of products of the reaction-phytanic acid and CoA (27, 28). Finally, the hydrolysis of ATP resets the transporter to inward open apo state.

## Materials and Methods

### Human ABCD3 expression and purification

The human ABCD3 (Uniprot ID: P28288) gene was synthesized by Genscript and cloned in pFastBac1 vector with a C-terminal 8xHis tag for baculovirus-mediated expression in *Spodoptera frugiperda* cell line *Sf9*. Bac-to-Bac™ Baculovirus Expression System (Thermo Fisher Scientific) protocols were followed to prepare recombinant baculovirus. Baculovirus multiplicity of infection (MOI) of 2 was used to infect the *Sf9* cells and cultured for 48 hours at 27°C. The cells were harvested by centrifugation and then resuspended in the Lysis buffer (50 mM Tris Cl, pH 7.5, 100 mM NaCl, 10 mM MgCl_2_, 10% glycerol) supplemented with 1X EDTA-free cOmplete protease inhibitor cocktail tablets (Sigma-Aldrich). Cells were lysed using sonication at 4°C and then dounce homogenized. 1% w/v lauryl maltose neopentyl glycol (LMNG) (Anatrace): 0.1% w/v cholesteryl hemisuccinate (CHS) (Anatrace) was added to the lysate to solubilize the membrane fraction overnight. Solubilized membrane was separated by centrifugation at 104,630 *xg* for 30 min. 5 mM imidazole added to the supernatant. HisPur™ Cobalt Resin (Thermo Fisher Scientific) was equilibrated with the Lysis buffer and 5 mM imidazole, and the protein in the supernatant was allowed to bind for 2 h on a shaking platform. The mixture was passed through a disposable gravity column and flowthrough was collected. The resin was washed with 10 mM imidazole, 0.02% glyco-diosgenin (GDN) (Anatrace) containing Lysis buffer. ABCD3 was eluted from the cobalt resin with 150 mM imidazole, 0.02% GDN containing lysis buffer. The protein was concentrated with Amicon Ultra-15 centrifugal filters (MilliporeSigma) and further purified using size exclusion chromatography (SEC) in SEC buffer (25 mM HEPES pH 7.5, 100 mM NaCl, 2 mM MgCl_2_, and 0.02% GDN). For ligand binding experiments and activity assays 0.005%:0.0005% LMNG:CHS was used instead of 0.2% GDN throughout the protocol.

### ATPase activity assay

ATPase assays were performed using malachite green based colorimetric ATPase/GTPase Activity Assay Kit (Sigma-Aldrich). ∼4.5 µg of purified ABCD3 in buffer 25 mM HEPES pH 7.5, 50 mM KCl, 0.005%:0.0005% LMNG:CHS was used to assess ATPase activity in a 40 µL reaction volume in triplicates. The reaction mixture above with 2 mM ATP/Mg^2+^ with or without substrate was incubated for 30 min at 37°C. The substrates phytanoyl-CoA and palmitoyl-CoA (Avanti Polar Lipids) when mentioned were added at 250 µM. The reaction was stopped by adding reagent from the assay kit and incubated for 30 min at room temperature. The colorimetric product corresponding to phosphate released was assessed by measuring absorbance at 620 mm using Tecan Spark® Multimode Microplate Reader.

### Cryo-EM sample preparation

For the apo structure, ABCD3 was used at ∼6 mg/mL. 3.5 µL of the sample was applied to Quantifoil R 1.2/1.3 300-mesh Au holey carbon grids. Fluorinated Fos-Choline-8 (Anatrace) at a final concentration of 1.5 mM was added directly on the grid and mixed with the sample. Mark IV Vitrobot (FEI) was used for sample vitrification on the grids at 8°C and 100% humidity. A blot force of −2 and blotting time between 3 and 7 s was used to remove excess sample from the grids. The grids were rapidly plunge frozen after the blotting in liquid ethane and stored in liquid nitrogen.

For the substrate bound structure, 0.5 mM phytanoyl-CoA was added to ABCD3 at a final concentration of ∼8 mg/mL. 3.5 µL of the sample was applied to freshly glow-discharged Quantifoil R 1.2/1.3 400-mesh Au holey carbon grids. The vitrification was done as described for the apo structure.

### Cryo-EM data acquisition

Imaging of ABCD3-apo and ABCD3-phytanoyl-CoA grids was done on a Titan Krios 300 keV. microscope. A defocus range of −0.8 to −2.5 was used at a total dose between 45 and 60 e/Å^2^. The Titan Krios was equipped with GATAN K3 direct electron detector and a post column energy filter with 20 eV slit width that was employed during data collection. SerialEM software was used for the data collection with automated data acquisition settings. Data acquisition details including the number of micrographs collected for each dataset, pixel size, dose, are listed in appendix, Table S1.

### Cryo-EM image processing and model building

Cryo-EM data processing was performed using CryoSPARC as summarized in appendix, Fig. S2 -S4 (29). Model building of the ABCD3 apo based on the 3.3 Å cryo-EM map was done in Coot (30). The real space refinement of the structure was performed in PHENIX to obtain the final model (31). ABCD3 apo model was used as a starting point for ABCD3-phytanoyl-CoA and was further fitted and refined in Coot and PHENIX, respectively. PhyCoA SMILES file based on the chemical structure was used to generate restraints in PHENIX eLBOW. The phytanoyl-CoA file from eLBOW was fitted in the cryo-EM density using coot. ABCD3 and phytanoyl-CoA together were refined in PHENIX. ChimeraX was used for visualization and figure preparation (32, 33). The above mentioned software were installed through SBGrid consortium (34)

### Phytanoyl-CoA binding analysis

For Microscale thermophoresis (MST) ligand binding experiments (35), purified ABCD3-8xHis protein, expressed in *Sf9* cells, was prepared as described and quantified by measuring absorbance at 280 nm using a NanoOne^C^ spectrophotometer (Thermo Fisher Scientific). The protein was labeled with Red-tris-NTA dye (NanoTemper) in SEC buffer for 30 minutes at room temperature. After labeling, the sample was centrifuged at 10,000 *xg* for 10 min to remove any protein aggregates. The resulting labeled protein was incubated at 100 nM with varying concentrations of phytanoyl-CoA for 30 minutes at room temperature. The protein-ligand mixtures were then transferred to premium capillaries (NanoTemper) for analysis using the Monolith NT.115 Pico instrument.

MST measurements were conducted in technical triplicate at 25°C with medium MST power (40%). Data were analyzed using MO Affinity Analysis software (v3.0.5) to calculate the dissociation constant (K_d_).

### Molecular dynamics simulation

All-atom molecular dynamics (MD) simulations were performed to investigate the protein and substrate dynamics within a lipid bilayer environment using the GROMACS simulation package (36) and the CHARMM36 force field (37). The MD simulations were initiated using the cryo-EM structure of the phytanoyl-CoA bound ABCD3 protein system. This system was embedded in a pre-equilibrated 1-palmitoyl-2-oleoyl-sn-glycero-3-phosphocholine (POPC) lipid bilayer, generated using the CHARMM-GUI web server (38) and oriented according to the PPM server (39). The system was solvated using the TIP3P water model and neutralized with Na^+^ and Cl^−^ ions to a final concentration of 150 mM, simulating physiological ionic strength. Following 5000 steps of steepest descent energy minimization, the system underwent a multi-stage equilibration protocol totaling 50 ns, during which positional restraints on protein heavy atoms and lipid phosphorus atoms were gradually released. An additional 50 ns of unrestrained NPT equilibration ensured thorough system relaxation prior to production simulations.

Three independent production MD simulations, each 500 ns in duration, were performed under the NPT ensemble, maintaining 303.15 K and 1.0 bar using the velocity-rescale thermostat and C-rescale semi-isotropic barostat, respectively. Electrostatic interactions were treated with the Particle Mesh Ewald (PME) method, while van der Waals interactions used a switched cutoff scheme. The LINCS algorithm constrained bonds involving hydrogen atoms, permitting a stable 4 fs integration timestep enabled by hydrogen mass repartitioning (HMR). Using these simulation parameters and settings, system equilibration was confirmed by analyzing membrane thickness and area per lipid over the final 50 ns of the NPT equilibration phase. Structural stability of both protein and ligand during the three independent production runs was subsequently evaluated via RMSD calculations of protein backbone Cα atoms and phytanoyl-CoA heavy atoms (Appendix Fig. S5).

## Data availability

The Electron Microscopy Data Bank (EMDB) accession codes for cryo-EM density maps are EMD-49388 for apo human ABCD3, and EMD-49386 for phytanoyl-CoA bound human ABCD3, respectively. The structural models are deposited in the Protein Data Bank (PDB) database under accession codes 9NGM (apo) and 9NGJ (phytanoyl-CoA bound).

## Contributions

M.G., N.K.K., R.M.S. designed research; M.G., N.K.K., D.J.S., S.G.B., M.S.D., A.M., I.E. performed research; M.G., N.K.K. contributed new reagents/analytic tools; M.G., N.K.K., D.J.S., M.S.D., A.M., S.G.B., I.E., R.M.S. analyzed data; M.G., R.M.S. obtained funding; and M.G., N.K.K., D.J.S., S.G.B., I.E., R.M.S. wrote the paper.

## Competing Interest Statement

Authors declare no competing interests.

## Acknowledgments

Research was supported by the National Institute on Aging (R00AG070271), and American Heart Association (835634) to M.G., and National Institute of General Medical Sciences (R01GM024485) to R.M.S. We thank Phoung Nguyen, Alisa Bowen for help with initial protein expression trials. We acknowledge David Bulkley, Glenn Gilbert, and Matt Harrington for advice with EM data collection carried out at the UCSF cryo-EM facility (supported by NIH grants S10OD020054, S10OD021741 and S10OD026881). A part of this work was performed at the Stanford-SLAC Cryo-EM Center (S2C2), which is supported by the National Institute of General Medical Sciences (1R24GM154186). The content is solely the responsibility of the authors and does not necessarily represent the official views of the National Institutes of Health. The authors would also like to thank the following S2C2 personnel for their invaluable support and assistance: Patrick Mitchell, Corey Hecksel, Megan Mayer, Lisa Dunn. We acknowledge the contributions of the OHSU Biophysics Shared Resource Core (Research Resource ID: RRID: SCR_022744) in facilitating this work.

## Notes

### Competing Interest Statement

The authors have declared no competing interest.

